# N,N-dimethylacetamide blocks inflammation-induced preterm birth and remediates maternal systemic immune responses

**DOI:** 10.1101/2025.01.16.633350

**Authors:** Sandra E. Reznik, Alexander Kashou, Daylan Ward, Steve M. Yellon

**Affiliations:** Department of Pharmaceutical Sciences, College of Pharmacy and Health Sciences, St. John’s University, 8000 Utopia Parkway, Queens, NY 11439; Departments of Pathology and Obstetrics and Gynecology and Women’s Health, Albert Einstein College of Medicine, Bronx, NY 10461; Longo Center for Perinatal Biology, Loma Linda University School of Medicine, Loma Linda, CA 92350

**Keywords:** cytokines, pregnancy, parturition, cervix

## Abstract

The common excipient, N,N-dimethylacetamide (DMA), prevents imminent endotoxin-induced preterm birth in mice. The present study hypothesized that DMA forestalls preterm birth to term (defined as day 18.5 or later) by attenuating bacterial endotoxin lipopolysaccharide (LPS)–induced maternal systemic inflammatory responses and cervix remodeling. Accordingly, LPS (i.p.) on day 15 postbreeding stimulated preterm delivery within 24 h while mice treated with DMA 2 h preceding and 9 h following LPS administration remained pregnant, comparable to saline and DMA controls, to deliver viable pups at term. Irrespective of LPS or DMA+LPS treatment, maternal plasma pro- and anti-inflammatory cytokines on day 15.5 (12 h post-LPS) increased 10-fold compared to baseline concentrations in controls. On day 16 of pregnancy, plasma concentrations of G-CSF and TNFα were reduced in the prepartum LPS+DMA group compared to those in postpartum mice given LPS. By day 18 of pregnancy, all cytokines returned to baseline - equivalent to low systemic levels throughout the study in saline and DMA controls that gave birth at term. In addition, maternal plasma progesterone declined within 12 h in prepartum LPS-treated mice to postpartum concentrations on day 16. Although a similar transient decrease occurred by 12 h in DMA+LPS mice, plasma progesterone returned to baseline concentrations in controls. Contemporaneously, the progression of prepartum cervix remodeling leading to preterm delivery was acutely forestalled by DMA without impeding birth at term. These findings support the hypothesis that DMA not only prevents inflammation-driven preterm birth, but rescues pregnancy for birth to occur at term. The results raise the possibility that maternal signals can forecast risk of preterm birth while selective suppression of systemic inflammation can mitigate adverse pregnancy outcomes.

**Summary:** DMA rescues pregnancy from inflammation-induced preterm birth through a progesterone-independent mechanism that involves multiple proinflammatory cytokines in maternal circulation.

Supported by NIH Grant 1R16GM145586 and the Ines Mandl Research Foundation

## Introduction

Parturition in mammals is a complex and varied process among species that requires coordinated functions by multiple organs. In women, obstetricians are challenged to answer when a baby will be born, whether at, before, or post-term. Upwards of 30% of women in the USA conclude pregnancy by cesarean delivery [1]. Preterm birth rates exceed 11% of all pregnancies in some countries, and remains the major cause of neonatal morbidity and mortality worldwide [2–6]. Preterm birth rates have not significantly improved over several decades despite major advances in medical technologies in patient care [7, 8]. A 2023 BMC review of the drug development pipeline to treat preterm birth and preterm labor identified no candidates with the potential to block premature delivery [9].

Assessment of maternal systemic indications of inflammation before term or in premature deliveries, whether spontaneous or induced, is challenging. Noteworthy is the finding that maternal systemic proinflammatory cytokines were elevated throughout uncomplicated pregnancies in women with a male versus female sex of the fetus, but relatively stable after 28 weeks to term [10]. Inflammation and infection are recognized risks for adverse pregnancy outcome, but availability of data for markers of systemic inflammation is limited [11, 12]. Lipopolysaccharide (LPS) is a widely used bacterial endotoxin in studies of inflammation-induced preterm birth. Although the toll-like receptor (TLR)-4 cellular mechanism of action is well-established [13–17], effects of LPS on the time course of maternal systemic inflammation or consequences of anti-inflammatory treatment to prevent preterm birth are not known. Moreover, the relationship of these immune responses and consequences of LPS treatment on cervix structure, the critical barrier that must remodel for birth to occur, is also unknown. Evidence from available peripartum cervix biopsies in women [18–21] and in studies of mice [22] indicate a local inflammatory process is associated with degradation of cross-linked collagen and fewer cell nuclei (CN) per area before birth. Thus, virtues of the well-characterized LPS-induced model for preterm birth in mice provides an opportunity to investigate the relationship of prepartum maternal systemic inflammatory markers for term and preterm birth.

N,N-dimethylacetamide (DMA), a widely used United States Food and Drug Administration (FDA) approved drug excipient, was discovered to have anti-inflammatory properties that block LPS-induced PTB in mice by a mechanism that inhibits the transcription factor NF-κB [23–26]. Furthermore, DMA suppresses cytokine secretion in various immune and placental cell phenotypes, including human trophoblast JEG-3 cells and placental explants [24], while also preventing degradation of the NF-κB inhibitory molecule, I kappa B alpha (IκBα) [25]. Additionally, DMA was found to be epigenetically active by binding bromodomains, such as Brd4 [27], which subsequently prevents activation of NF-kB [28]. A closely related analog to DMA, dimethylformamide (DMF), also used in the pharmaceutical industry, was found to suppresses IL-8 and IL-6 secretion from LPS-challenged human placental trophoblast HTR-8/SVneo cells and to delay preterm birth in LPS-induced C57Bl/6 mice [29]. With the potential anti-inflammatory therapeutic value of DMA, the aim of this investigation was to determine if DMA suppressed the time course of a maternal systemic inflammatory response to LPS to forestall prepartum cervix remodeling and delay birth to term.

## Methods

### Animals and experimental design

All methods related to animal experiments are reported in accordance with ARRIVE guidelines. Adult nulliparous, timed pregnant CD-1 mice were purchased from Charles River Laboratories (Hollister, CA) and individually housed in the Loma Linda University vivarium in controlled conditions with 12-h lights on, at 7 am PST. All procedures were conducted under an Institutional Animal Care and Use Committee-approved protocol that conformed to NIH Office of Laboratory Animal Welfare guidelines. D15 mice were divided into four groups, and received the following administrations intraperitoneally (i.p.): 1) saline (0.1 mL) at 7 am, 9 am and 6 pm (saline controls, 0.1 M phosphate buffered saline); 2) DMA (20 mmol/kg, 0.1 ml, ThermoFisher Cat #390670010) at 7 am and 6 pm and saline at 9 am (DMA controls); 3) the TLR-4 ligand lipopolysaccharide (LPS, 2 mg/kg = 80 µg/0.1 mL, serotype O55:B5, Sigma Aldrich, St. Louis, MO) at 9 am and saline at 7 am and 6 pm (LPS controls); and 4) LPS at 9 am and DMA at 7 am and 6 pm (DMA+LPS mice). Given variations in response to LPS, the dose used in the present study (70 and 80 µg) was based upon previous findings [30] and an initial pilot study using a dose range of 50-100 µg LPS/0.1 ml saline i.p. (n=3/group) to optimally induce 100% preterm birth with no signs of maternal morbidity or mortality. Mice in each group were euthanized by CO_2_ asphyxiation on D15.5, D16 and D18 (prepartum); others were allowed to proceed to delivery. Birth was defined as at least 1 pup ex-utero.

### Progesterone ELISA and multiplex cytokine immunoassay

Blood was collected into 0.1 M EDTA-treated tubes immediately after each mouse was euthanized. Plasma was harvested and frozen at -80°C until thawed and duplicate aliquots were assayed for progesterone (0.04 ml) by Diagnostics (MN, USA, using 3 Enzo mouse kit, Cat# ADI-900-011). Average assay sensitivity was 0.2 ng/ml, while inter-assay coefficients of variability were <5%. In addition, duplicate aliquots (0.025 ml) were assayed using 3 Custom BioPlex kits and Bio-Plex 200 system according to kit specifications. These cytokines reflect a survey of pro- and anti-inflammatory functions that mediate effects of infection and are prominent in the preterm birth process and/or the NF-κB signaling pathway. Range of assay sensitivity (pg/ml) was for granulocyte colony stimulating factor (G-CSF) 4.72-4.93, granulocyte-macrophage colony-stimulating factor (GM-CSF) 2.05-4.71, interferon gamma (IFNγ) 1.32-1.49, Interleukin-1 beta (IL-1β) 2.18-2.19, IL-4 0.66-0.71, IL-6 0.96-1.13, IL-10 3.31-4.33, IL-12 4.27-4.60, monocyte chemoattractant protein-1 (MCP-1) 10.00-15.03, and tumor necrosis factor alpha (TNFα) 3.50-4.55. Coefficients of variation were for intra-assay duplicates 5.0, 1.9, 1.6, 31.6, 13.3, 4.7, 1.3, 4.8, 6.2, 2.4, and inter-assay kits 7.0, 35.7, 9.9, 48.1, 11.7, 12.8, 12.7, 14.5, 8.8,17.7, respectively.

### Assessment of cross-linked collagen and cell nuclei density in peripartum cervix

Postmortem, the cervix was excised, immersion fixed, and adjacent sections stained with either picrosirius red (PSR) to evaluate collagen cross-linking, an indication of remodeling, or methyl green to count cell nuclei. As previously described, the normalization of OD/cell nuclei density has served as an index of inflammation (edema/cellular hypertrophy) [31–34]. For this study, Visiopharm software was used to develop a novel automated algorithm for cell nuclei counting. The validation of this approach is detailed in the Supplement. A survey of 6-8 photomicrographs in regions of interest (ROIs) within duplicate 10 μm sections from the ectocervix and endocervix stroma from each cervix was analyzed under circularly polarized light. The optical density of picrosirius red stain birefringence under polarized light is specific for collagen and inversely proportional to the density of cross-linked fibrillary structure [35, 36]. This analysis of collagen staining is the only *in situ* approach to specifically assess extracellular cross-linked collagen in the intact cervix stroma, has proven useful for similar analyses in other fixed tissues [22], and correlated with biochemical and electron microscopic analysis of fiber length and diameter in cervix stroma [31, 35].

### Statistical analyses

Parametric statistics were rigorously applied to analyze each dataset (progesterone, individual cytokines, cell nuclei and optical density). Data were initially analyzed by Grubb’s and Levene’s tests, respectively for outliers and variance of data. Subsequently, for these qualified datasets with equal variance homogeneity, 1-way ANOVA with Tukey’s test for individual comparisons was used within each of the 4 groups across days of the study. In addition, among treatment groups on the same day of study, the 1-way ANOVA with Tukey’s test for individual comparisons was used. For some datasets, a 2-way ANOVA with Tukey’s test best suited the experimental design because mice injected with PBS (Con) served as controls for LPS treatment while mice administered DMA served as controls for the DMA+LPS treatment. The use of 2-way ANOVA with Sidak’s test reduced extraneous multiple comparisons to improve sensitivity of the statistical analysis. However, not all data had equal homogeneity of variance. Consequently, data were log transformed and, if variance were equal, analyzed as described above. When original and log transformed data both displayed unequal variance homogeneity, non-parametric Kruskal-Wallis with Dunn’s test was utilized on the original data for the same comparisons as the 1-way ANOVA.

## Results

### DMA prevents LPS-induced preterm birth and forestalls the decline in maternal systemic progesterone

Controls given saline (Con) or DMA gave birth at term, between days 18.5 and 21 (Fig. 1 panel a). By contrast, nine out of 13 mice (69.2%) treated with LPS delivered preterm, with birth of at least 1 pup within 24 h, by day 16 postbreeding. Importantly, DMA+LPS-treated mice did not deliver until, on average, term in a time line comparable to that of controls. Moreover, pups that were born at term in to dams in the control, DMA and DMA+LPS groups as well as in 4 LPS treated mice that did not deliver preterm, had milk in their stomachs, an indication of sufficient postpartum vitality to nurse.

**Figure 1.**
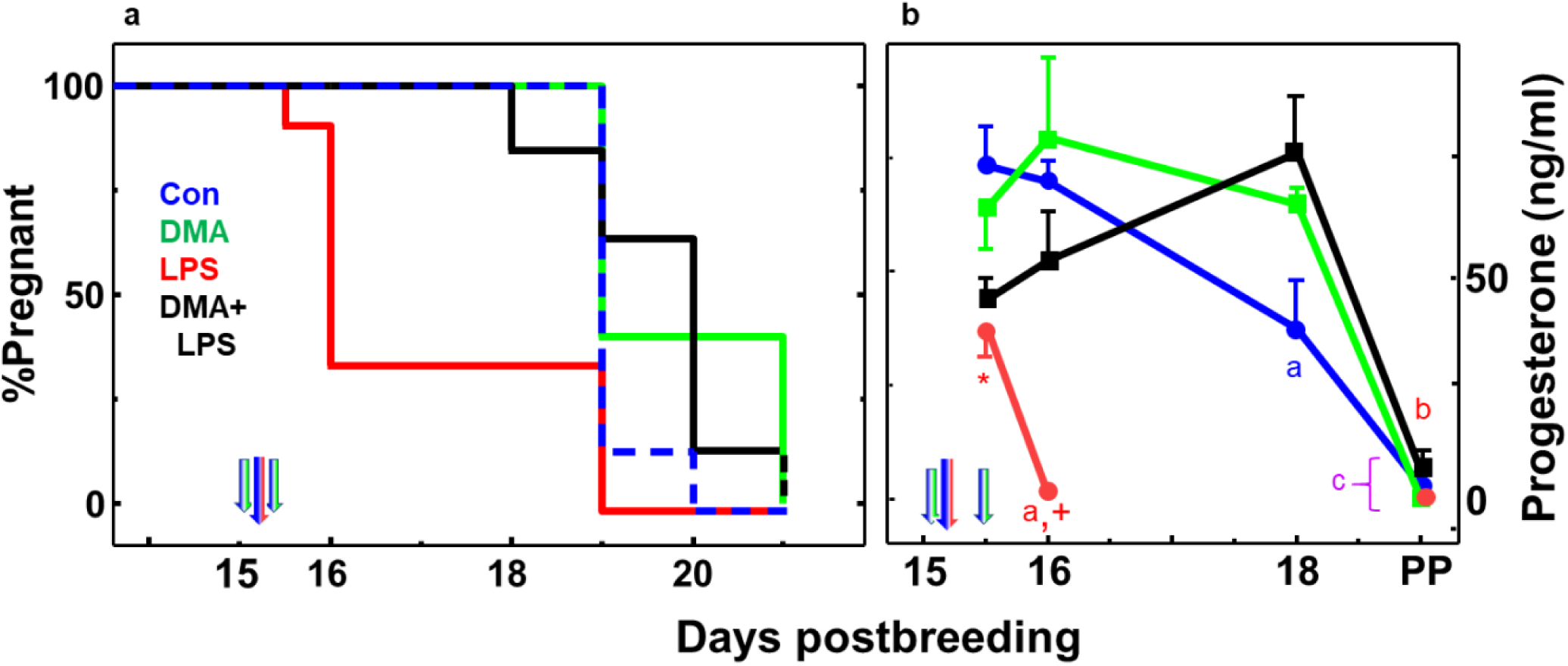
Panel a. DMA prevents LPS-induced preterm birth. Kaplan-Meier plot of birth in mice injected on day 15 of pregnancy (D15) with saline or DMA at 0700 h and 1800 h saline (blue/green arrows), as well as saline or LPS at 0900 h (blue/red arrow). Thus, the 4 groups were **Control** (**Con**, blue dashed line; saline only; n= 20), **LPS** (red line; saline+LPS; n= 18)**, DMA** (green line; 2 doses of DMA in place of saline+saline in place of LPS; n= 15), or **DMA+LPS** (black line; 2 doses of DMA, 2 h before and 9 h after LPS; n= 23). See Methods for study detail. **Panel b. DMA rescues LPS-induced suppression of progesterone in maternal circulation**. Plasma progesterone concentrations in same groups of mice described in panel a (mean ± SE, n= 3-8/group/time). SE for some time points are covered by area of symbol. All mice in the D15.5 groups were prepartum, 12 h after mice were treated with saline (Con, blue half arrow) or LPS (red half arrow) to induce preterm birth. By day 16, 9 of 13 mice in the LPS group (red line) were postpartum (PP), while 4 gave birth at term (D19). Mice in all other groups birthed at term (average day 19.5 postbreeding). Briefly, statistical analyses included Grubb’s test to exclude outliers and Levene’s median test for homogeneity of variance. Afterwards, 1-way ANOVA with Tukey’s test for multiple comparisons was performed on log transformed data. p<0.05 was indicated compared to **^a^** all previous days for same treatment group and **^b^** D15.5 same group. With respect to same day postbreeding comparisons, **\*** D15.5 LPS versus Con, and **^+^** D16 LPS versus all other groups. See Methods for ELISA assay and statistical analyses details.

As expected, mortality among the pups delivered by Con mice was minimal (0.7%, 2 of 269 pups were stillborn). Morbidity and mortality among the DMA control pups was 12%, 23 of 187 pups – mostly stillborn some resorbtion implantation sites, and one litter cannabilized. In the LPS group that delivered preterm, most pups were cannabilized so morbidity and mortality can not be accurately assessed.

Finally, morbidity of pups in dams treated with DMA+LPS was 30%, (104 of 344). No maternal mortality occurred in Con, DMA or LPS groups, but two dams died in the DMA+LPS group within 2 days of treatment (7%, 2 of 28).

For progesterone, plasma concentrations in pregnant controls given saline or DMA alone were equivalently elevated for the duration of pregnancy (Fig. 1 panel b, blue or green lines, respectively). Within 12 h of LPS injection, systemic progesterone was suppressed in dams in the LPS and DMA+LPS groups on day 15.5 compared to saline and DMA controls, respectively. By day 16, maternal progesterone in 9 postpartum LPS-treated mice had fallen to baseline. However, on day 16 and continuing to day 18 of pregnancy, plasma progesterone was similarly elevated among controls (saline and DMA groups), as well as in dams given DMA+LPS on day 15. With birth at term (average day 19.5 postbreeding) and in 4 LPS-treated mice whose pregnancy progressed to term, plasma progesterone declined to baseline.

### DMA remediates LPS-induced responses by proinflammatory cytokines in maternal circulation and sustains pregnancy

Compared to controls given saline only or DMA (both received saline at 9 am instead of endotoxin), LPS induced a greater than 10-fold increase in the prepartum maternal circulation of all ten cytokines within 12 h irrespective of DMA treatment (Figs. 2 and 3). Moreover, while all cytokines continued to increase in the next 12 h in the LPS mice (day 16 postbreeding 24 h after LPS), remarkably, the change in cytokine levels reversed direction in all cases in DMA+LPS mice (Figures 2 and 3). With the exception of GM-CSF, which, interestingly, rebounded after D18 (Figure 3), cytokines in DMA+LPS mice dropped to levels in controls given saline only or DMA by D18 and remained there until parturition. While DMA clearly affected all ten cytokines investigated, based on its ability to reverse the direction of cytokine expression between D15.5 and D16, the difference in cytokine concentrations between LPS and DMA+LPS dams at D16 was statistically significant for only two cytokines tested: G-CSF and TNFα (Figure 2, top panels).

**Figure 2.**
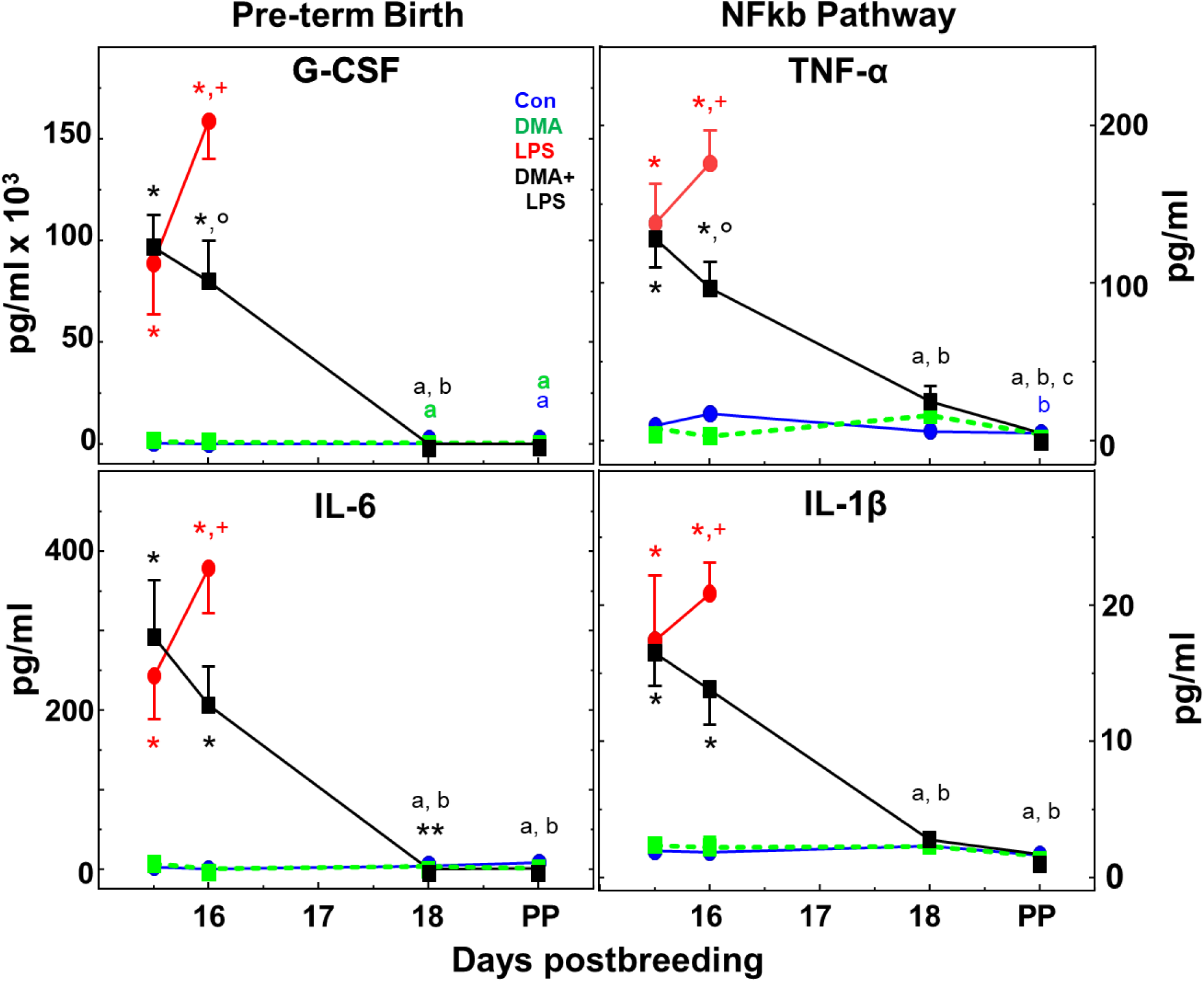
DMA mitigates the LPS-induced rise in proinflammatory cytokines in maternal circulation. Data are mean ± SE (average of n=3-9/group/day postbreeding). Group designations are the same as in the Figure 1. Area of symbol may cover the SE. As detailed in the Methods for Statistical analyses, this dataset was qualified by passing Grubb’s and Levene’s tests with log transformation and ANOVAs performed. p<0.05 was indicated by 1-way analysis within groups across each day versus ^a^D15.5, ^b^D16, and ^c^D18. 1-way analysis was also used within each day across groups, specifically *Con and DMA versus LPS and DMA+LPS, **D18 DMA+LPS versus Con only (all at or near assay baseline sensitivity), as well as ^+^LPS D16PP versus all other PP Term. Additionally, a 2-way ANOVA with Sidak’s Test was performed on treatment groups LPS and DMA+LPS for days 15.5 and 16. p<0.05 for **°** D16PP LPS versus D16 DMA+LPS. Colors of statistical symbols correlate with group comparison.

**Figure 3.**
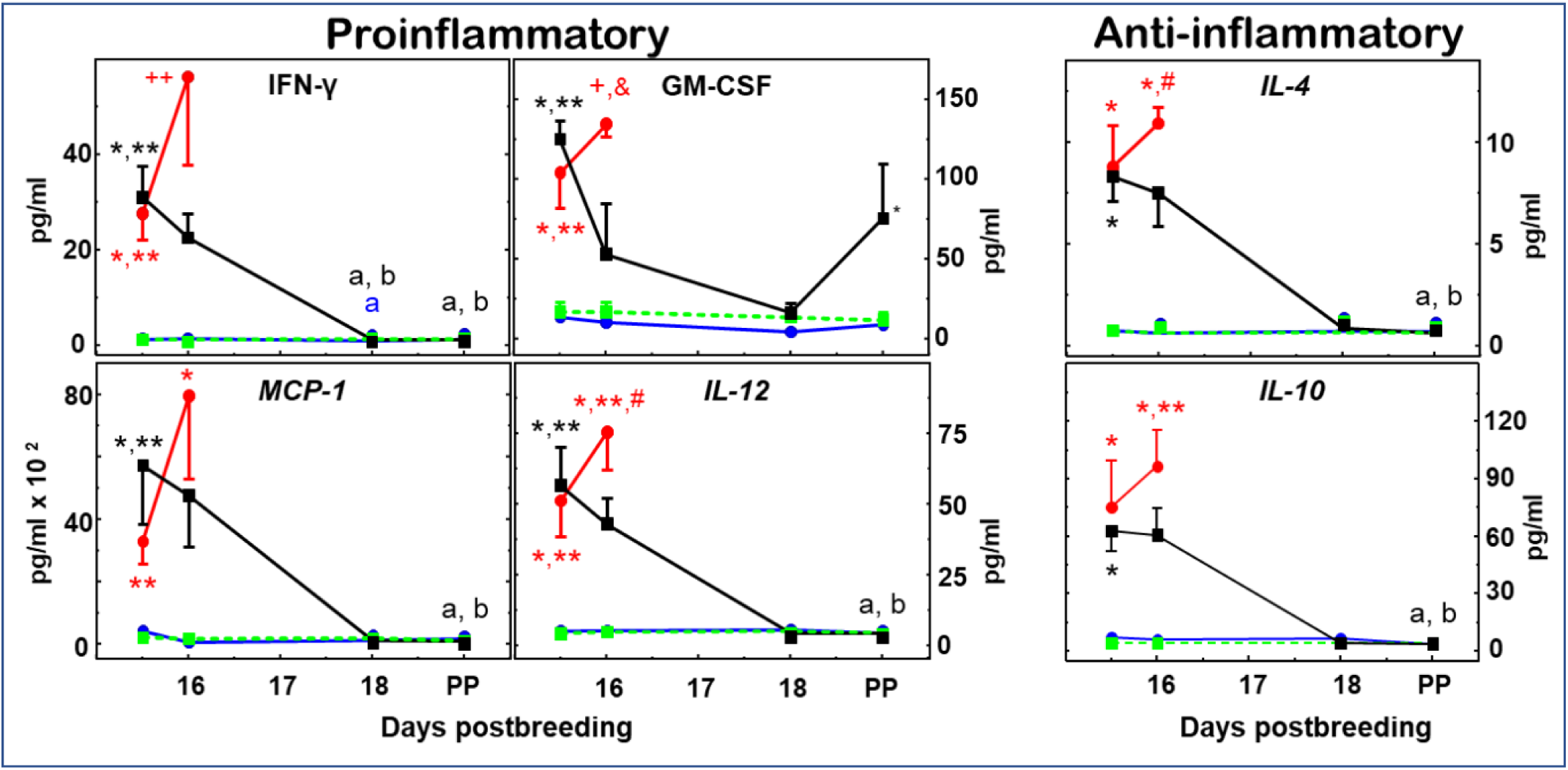
Proinflammatory and anti-inflammatory cytokines induced by LPS in maternal circulation are reduced to that in controls within 2 days by DMA. Group designations are the same as that in Figure legends 1 and 2. Data are the mean ± SE (average for Con and DMA groups n=4, range 3--9/group/day postbreeding, for LPS and DMA+LPS groups average n=7, range 3-9). IFNγ data passed Grubb’s and Levene’s tests on untransformed data and statistical analysis done as in Fig. 2. p<0.05 for 1-way analysis within groups across each day versus ^a^D15.5 and ^b^D16. Within each day across groups, 1-way indicated p<0.05 versus *Con,**DMA, and ^++^ all other PP term. For GM-CSF, parametric analysis was performed on log -transformed data while for MCP-1, IL-12, IL-4, and IL-10, both untransformed and log-transformed data had unequal homogeneity of variance (Levene’s test p<0.05), which necessitated use of nonparametric Kruskal-Wallis and Dunn’s tests. p<0.05 indicates versus ^a^D15.5 and ^b^D16 within groups across days. While across groups within day, p<0.05 indicates *Con,**DMA, ^#^Con and DMA+LPS PP term, ^+^Con and DMA PP term, and ^&^ D16PP LPS versus all other D16.

Figure 3 findings indicate that concentrations of the other four pro-inflammatory (IFN-γ, GM-CSF, MCP-1 and IL-12) and two anti-inflammatory cytokines (IL-4 and IL-10) also increased compared to controls on days 15.5 and 16 (Con and DMA groups), but differences in maternal circulation did not reach statistical significance in the DMA+LPS versus LPS groups on day 16. With the exception of GM-CSF, other cytokines were at or near baseline by the day of birth at term and comparable to that in postpartum controls. Importantly, for saline- or DMA-treated mice in controls, cytokines were at or near baseline throughout the study. Moreover in the 4 LPS-treated mice that gave birth at term, all plasma cytokines in the maternal circulation were at levels equivalent to those in controls at term. Thus for the cytokines in this study, the findings highlight the absence of maternal systemic inflammation in the process of parturition at term as opposed to preterm birth.

Two approaches were taken to assess remodeling of the prepartum cervix associated with effects of treatment leading up to term or preterm birth. Density of cell nuclei, an assessment of cervix remodeling due to edema or cellular hypertrophy, was counted as in previous publications and data were used to validate an automated image analysis algorithm to enumerate cell numbers/area in the same photomicrographs (details in Methods and the supplement). The decline in the cell nuclei density found in prepartm controls given saline by D18 postbreeding was not seen in DMA-treated mice (Fig. 4, top panel). However, by D16 in LPS-treated mice, the fall in density of cell nuclei in the cervix occurred in the groups that gave birth preterm and by D19 in the four mice that delivered at term. In the DMA+LPS-treated group, the effects of LPS were blocked such that cell nuclei density was unchanged on D16, but reduced on D18 and at term-comparable to that in controls.

**Figure 4.**
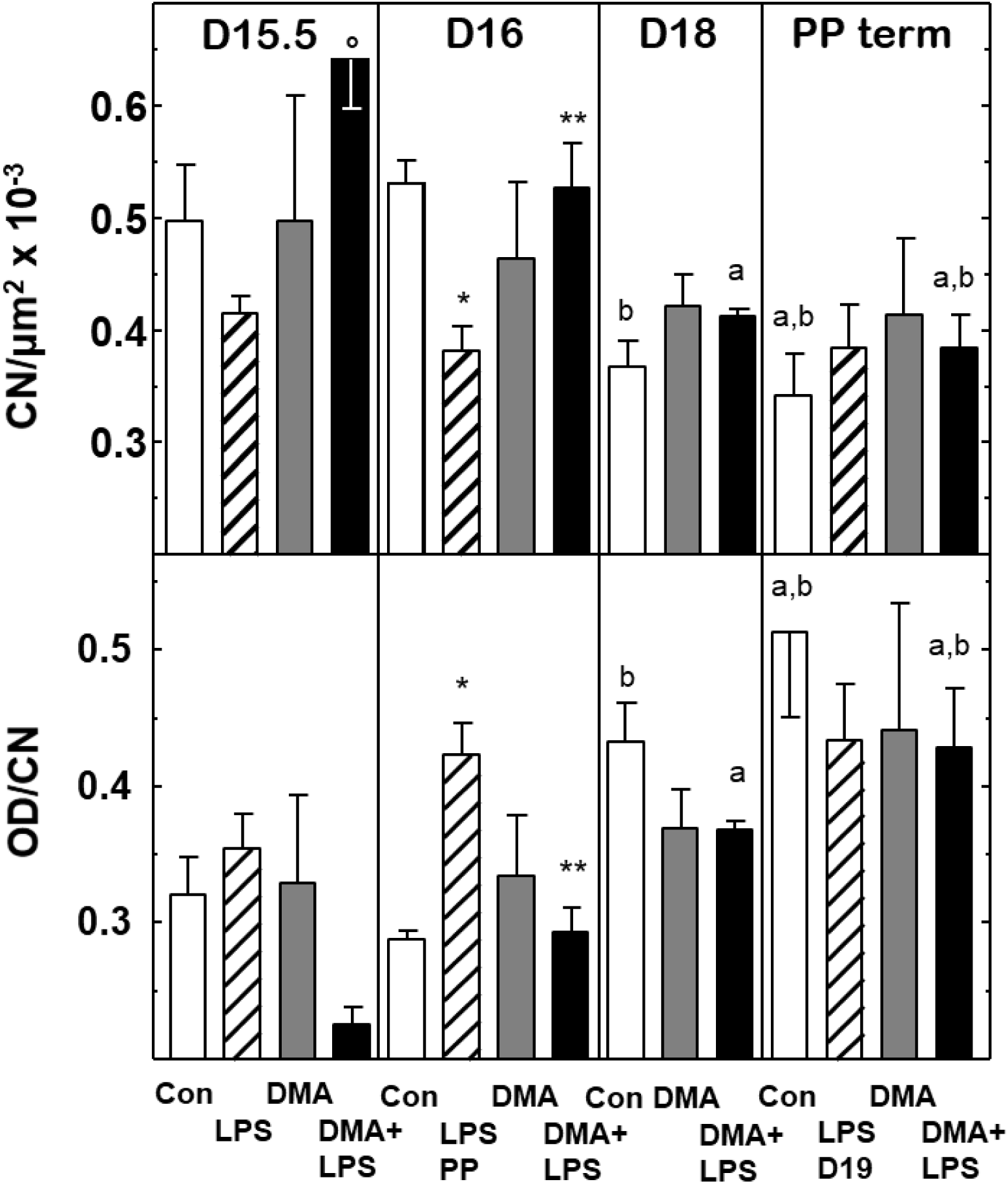
Panel a. Degradation of cross-linked collagen, as indicated by the inverse of optical density/cell nuclei (OD/CN), is reduced in the prepartum cervix stroma in saline controls, but blunted by DMA treatment before term birth. Cell nuclei density in ROIs from adjacent sections from the same mice. **Panel b.** Density of cross-linking collagen per CN and were assessed in sections of cervix stroma from mice at on the same days postbreeding (4 groups, n=3-7 mice/day) as previously described in Figure 1 legend and detailed in Methods. Statistical analyses followed the approach of Fig. 1, except ANOVAs were run on untransformed data. Within each group across all days, 1-way analysis p<0.05 indicated **^a^**D15.5 and **^b^** D16. Within each day across all groups, 1-way analysis p<0.05 indicated **\***Con and **\*\***LPS. To focus on relevant comparisons during cervix remodeling, a 2-way ANOVA with Tukey’s Test was performed on all groups for days 15.5 and PP. p<0.05 indicates °D15.5 D+L v LPS for CN.

Degradation of cross-linked collagen structure as the cervix remodels, reflected by increasing optical density of PSR birefringence adjusted to cell nuclei density (OD/CN, Fig. 4, bottom panel), was found on D18 postbreeding in prepartum controls and postpartum at term. Decreased cross-linked collagen occurred in postpartum LPS-treated mice by D16. While cross-linked degradation was evident in postpartum term groups of controls, as well as LPS- and DMA+LPS-treated mice, the decline in cross-linked collagen that was found in these groups did not occur in DMA-treated mice. In controls given saline, OD/CN increased in the preterm cervix stroma of mice by D18 compared to D15.5 postbreeding. DMA treatment blunted this rise such that there was no prepartum change in collagen cross-linking (p>0.05). In mice injected with LPS, OD/CN was unchanged 12 h after treatment, but collagen crosslinking was reduced within 24 h in coincidence with delivery on day 16 postbreeding compared to term birth in other groups. Cross-linked collagen was also reduced by DMA in prepartum mice given LPS, but not the next day when DMA+LPS gave birth at term. Though timing differed, prepartum CN density was reduced in mice both on the day before and the day of birth (Fig. 4, panel b; D15.5 LPS and D18 Con versus D15.5 Con; PP Con and PP LPS D16 versus D15.5 Con). Optical density and density of cell nuclei were not significantly different among all groups postpartum at term, as well as LPS PP D16 that delivered preterm. Thus, DMA treatment on day 15 postbreeding forestalled prepartum cervix ripening, but did not interfere with processes that result in parturition at term.

## Discussion

The findings indicate that DMA prevented mice from inflammation-induced preterm birth such that pups delivered at term. While LPS-induced preterm birth does not represent all causes of birth before 37 weeks of pregnancy, this model reflects a common cause of premature birth that may underestimate asymptomatic systemic or local inflammation during pregnancy. Comparison of systemic versus intra-amnionic LPS administration in a recent report directly found merit to both approaches in murine studies [37]. Like previous acute effects of DMA in LPS-treated C57Bl/6 mice [23], present findings are consistent with little change in proinflammatory cytokines during the weeks preceding birth at term in women [10]. Moreover, these results for the first time indicate that DMA treatments on D15 did not affect the mechanism of term parturition or the capability of dams and pups to nurse. Systemic increases in all pro- and anti-inflammatory cytokines studied were robust and unimpaired by 12 h after LPS treatment. However, within 24 h (D16), responses by two proinflammatory cytokines, G-CSF and TNFα, in DMA+LPS-treated mice were reduced compared to those given LPS that delivered preterm. Thereafter, within 72 h, the DMA treatment of LPS-challenged mice suppressed both pro- and anti-inflammatory cytokines in circulation to baseline concentrations found in controls on D18. In particular, DMA suppressed MCP-1 (CCL-2), likely a critical effect, as this mediator is expressed in trophoblasts, decidua and myometrium, recruits immune cells and is increased in preterm labor [38]. Interestingly, DMA also suppressed IL-10 after D16. As IL-10 has both pro-inflammatory and anti-inflammatory properties [39], the DMA-mediated decrease in IL-10 levels observed here may be an anti-inflammatory effect. Importantly, no evidence of maternal systemic inflammation was found in saline or DMA-only treated controls throughout this period of pregnancy leading up to term. Thus, preterm DMA treatment did not affect the mechanism for parturition at term. Collectively, these data suggest that DMA blocks endotoxin-mediated preterm birth through a mechanism of action that may not involve dampening of anti-inflammatory cytokines or interference with acute host innate immune defenses. While beyond the scope of the current study, further investigation of the pharmacokinetics and route of administration of DMA on prepartum systemic, as well as local biomarkers of inflammation in relevant reproductive tissues are needed to further differentiate mechanisms of preterm compared to term birth.

As the potential reproductive toxicity of DMA in this model will be explored in future planned studies, pups were not maintained for a long period but were allowed to nurse for only 4 h. Of note, animal studies that have found potential reproductive toxicity associated with DMA had entirely different dosing schedules than the current study and included doses that were administered much earlier in pregnancy. In this study, DMA is administered late in gestation, once organogenesis is complete. The human equivalent of the DMA dose used in this study, using the converstion factor accounting for differences in body surface area [40] is approximately 10 mL of DMA (130 mg/kg) and could very easily be administered intravenously. Of course, concerns related to teratogenicity of DMA and, in fact, any putative approach to rescue pregnancy from preterm labor persist and have hindered the investigation of all types of pharmacotherapy to prevent premature birth. One important advance in the field is the development of vaginal nanoformulations that can deliver their cargo directly to the uterus and cervix, thereby decreasing exposure of the fetuses to a drug [41, 42]. In a line of investigation beyond the scope of the current study, a nanoformulation of DMA for local treatment may hold promise to reduce local inflammation associated with premature cervix remodeling using an already FDA-approved drug excipient at far below allowable daily doses.

The effects of LPS to induce preterm birth in this and other studies [43, 44] may result from pathways that are mediated by inflammation or impaired progesterone production or both. Within 12 h of LPS treatment, progesterone concentrations were equivalently reduced in prepartum mice irrespective of DMA treatment (Fig. 1 panel b). However, as noted in Methods (pilot and initial study), four mice did not respond to LPS and delivered at term. The potential to study other lower doses of LPS that induce a lower incidence of preterm birth may provide an opportunity to determine what systemic biomarkers of inflammation may predict risk of preterm birth. The blunted response to endotoxin in a small subset of mice underscores an advantage of studying immune responses in an outbred animal model - genetic diversity in molecular mechanisms may contribute to inflammatory drive to reduce from term the duration of gestation [45]. These findings raise the possibility that differential maternal responses to inflammatory stimuli could diminish critical components of the pathway that sustains pregnancy.

Findings clearly indicate that DMA did not by itself affect systemic progesterone concentrations nor block the effects of LPS to suppress plasma progesterone on day 15.5 postbreeding (Fig. 1 panel b). Characteristics that distinguished preterm birth in LPS-treated mice from prepartum DMA+LPS mice on day 16 were sustained plasma progesterone and reduced systemic G-CSF and TNFα. Moreover, DMA effects persisted because progesterone remained at peak, while concentrations of all cytokines were reduced to baseline in circulation within 2 days (by day 18). Thus, treatments with DMA on only day 15 postbreeding did not impact the mechanism for term parturition and sufficiently arrested an inflammatory drive for birth to occur at term.

Though premature to speculate whether DMA, indirectly or directly, affects the hypothalamic-ovarian axis to regulate responsiveness of reproductive tissues to progesterone in maternal circulation, it is conceivable that LPS via a TLR4-mediated inhibition of transcription factor NF-κB may affect systemic progesterone concentrations and response to progesterone by reproductive tissue. Enhanced progesterone in these mice 72 h after DMA+LPS administration may reflect the long-term consequences of the mechanism of DMA action to inhibit NF-κB [23–25]. DMA is predicted to decrease the synthesis of cyclooxygenases I and II (COX I and II) via its inhibitory effect on NF-κB. Inhibition of COX decreases prostaglandin F2α (PGF2α) synthesis and PGF2α synthesis leads to decreased production of progesterone in rodents [46]. Hence, DMA most likely prevents the high levels of PGF2α that are induced by LPS and thereby prevents the fall in progesterone that is observed in the LPS control mice. On the other hand, the effects of DMA may be independent of progesterone action since LPS and DMA+LPS treatment suppressed progesterone to an equivalent extent on D15.5. There may be long-term consequences of DMA mediated rescue of pregnancy with significant increases in systemic progesterone to concentrations significantly above those seen in controls. Perhaps this sustained elevated progesterone in DMA+LPS vs Con D18 contributed to pup mortality in the DMA+LPS PP group.Timing of the relatively small transient fall and recovery of plasma progesterone within 24 h of treatment in the DMA+LPS group is unlikely to explain the rescue of pregnancy compared to preterm birth after LPS. Alternatively, DMA may have a direct primary action to regulate progesterone in maternal circulation. Although precedence in the literature to support this contention is lacking, maternal progesterone concentrations in DMA-treated pregnant mice were the same as those in controls in the present study. Although DMA did not block the transitory decline in plasma progesterone in the DMA+LPS group 12 h after LPS, a rebound effect on plasma progesterone concentrations was found in the D18 DMA+LPS group versus controls.

Previous studies that used optical density of picrosirius red birefringence [35] and cell nuclei density [47] in the prepartum ectocervix and endocervix stroma of mice at term implicate these morphological characteristics are indicative of reduced collagen cross-linking, cellular hypertrophy, and edema. As an early part of the inflammatory process of ripening that occurs well-before labor and birth at term [18, 22, 31, 47–49], the present findings replicate past results in controls that found increasing OD/CN (decreasing collagen cross-linking) as well as decreasing CN density (increasing edema) from prepartum day 15 to 18, the 4 days before the day of birth at term. These findings highlight a counterintuitive misperception from use of the term ‘uterine cervix’. In actuality, the cervix is predominantly a connective tissue structure that is distinct from the smooth muscle, glands, and endometrium of the uterus [22, 50]. Unlike the diversity among species in morphology and function of the uterus, the cervix is highly conserved among mammals with a singular canal surrounded by similar phenotypes of cells, and a luminal epithelial zone that produces a unique mucus layer barrier [51].

Indeed, the cervix may function as an external maternal-fetal interface compared to other organs within the uterus which have become appreciated as an internal maternal fetal interface. The present study shows that DMA blunts 1) prepartum degradation of collagen cross-linking and 2) prepartum as well as LPS-induced cervix stroma edema reflected by reduced CN density (Fig. 4 top panel) without preventing normal parturition at term. Collectively, present findings raise the possibility that the inflammatory process induced prematurely by LPS and remediated by DMA may be distinct from the prepartum mechanism that remodels the cervix before active labor and parturition at term. These prospects require further investigation.

Although the pathways that lead to preterm versus term birth are likely to differ, a final common mechanism is proposed that converges upon prepartum local inflammation of the cervix for remodeling that depends upon loss of response to progesterone. Cytoarchitectural changes that are indicative of remodeling of the prepartum cervix before both inflammation-induced preterm birth and term birth are relevant because systemic progesterone concentrations are at or near peak and well before labor or birth [22]. Thus, across mammalian species, the loss of response to progesterone rather than a decrease in systemic progesterone appears to be a common feature associated with prepartum cervix ripening. Moreover, different mechanisms may differentiate preterm from term birth, but similarly remodel the cervix - whereas the former occurs with high cytokines in maternal circulation (biomarkers of systemic inflammation), the latter happens with evidence of only local inflammation in the cervix.

Of potential clinical relevance, pathogenic infection-related challenges during pregnancy including bacterial vaginosis and chorioamnionitis, are recognized major causes for preterm birth in women [52–54]. Based upon present findings that DMA arrests inflammation-induced preterm birth, possibly through differential suppression of particular proinflammatory cytokines, diagnostic and therapeutic approaches may be conceived to identify patients at risk for premature labor and the subset that progresses to the early conclusion of pregnancy. Focus on G-CSF and TNFα is warranted because maternal concentrations of these two cytokines were reduced within 24 h in the D16 DMA+LPS group compared to LPS-treated mice that delivered preterm. These cytokines have been implicated in the mechanism of preterm birth in women and mouse models of infection during pregnancy [55–59]. Even with known limitation in animal models [44], it is conceivable that assessment of certain proinflammatory signals in the maternal circulation in more advanced mammals may forecast risk for preterm labor or progression to premature birth. From a translational perspective, the opportunity to selectively suppress these cytokines or adaptive T cell immune responses to infection, without interfering with innate immune responses or term parturition, could sustain tolerogenic processes that maintain maternal well-being and typical duration of pregnancy. Ultimately, findings in this research model indicate that systemic inflammatory biomarkers in maternal circulation are not associated with parturition at term. By contrast, with endotoxin-induced preterm birth, systemic increases in cytokines precede prepartum cervical remodeling, while anti-inflammatory treatment with DMA restores plasma cytokines to pre-LPS concentrations, thus preventing inflammation in the cervix stroma and the continuation of pregnancy to term.

## Acknowledgments

Images were acquired for analyses in the California Tumor Tissue Registry Center in the Department of Pathology and Human Anatomy and the Advanced Imaging and Microscopy Core at the Loma Linda University School of Medicine. This project was in part supported by NIH HD054931, the LLU Department of Pediatrics Research Fund, NSF MRI-DBI 0923559 and NIH 1R16GM145586 and the Ines Mandl Research Foundation. In addition to the help of Anne C. Heuerman with data acquisition and initial analyses, the efforts by Ashley Thompson in preparation for experimental procedures, data analyses and graphics, as well as manuscript revisions were appreciated. The expertise Dr. Brit Boehmer was appreciated for development of the Visiopharm software algorithm.

## Author’s roles and agreed order of list

**Sandra E. Reznik** conceived and collaborated in design of study, and equally co-wrote the manuscript.

**Alexander Kashou** performed cytokine data analyses and assisted in the draft and manuscript revisions.

**Daylan Ward** organized study procedures, performed BioPlex assays, data analyses, graphs, and manuscript edits.

**Steve M. Yellon** conceived, designed, and supervised experiments, and equally co-wrote the manuscript.

## Data availability

Data are provided within the manuscript and the Supplementary information file.

