## Supplemental Figure 1 and Supplemental Table 1 for "N,N-dimethylacetamide blocks inflammation-induced preterm birth and remediates maternal systemic immune responses"

Supplement

Estimates of CN density came from manual counts of methyl green-counter-stained CN in a survey of 6-8 snapshots from scans of 1-2 longitudinal (sagittal) sections of the cervix with adjacent vaginal and uterine tissue [22, 30, 32, 33]. CN were identified in each snapshot (0.625 x10^5^ μm^2^/ROI) and recorded using NIH Image J. A random selection of ROIs with diverse counts were then checked by multiple individuals (students or staff) that were comparably-trained to identify distinct size, shape, and intensity of histochemically stained objects in the stroma of ectocervix and endocervix subregions. Cervix subepithelium, lumen, and vascular structures were excluded from consideration. A portion of a dataset of untreated controls were taken from a previous study [34, 36]. Manual CN counts were done on 6 snapshots/cervix, as described before, from 2 methyl green-stained scans/mouse at prepartum days 15.5, 16.0, 18.0; n= 7, 6, and 5, respectively, as well as postpartum day of birth (average of day 19.5 postbreeding; n=4). An unbiased standardized approach to recheck CN counts was to randomly select low, median, and high counts from individual mice across various days of pregnancy and background stain variations, irrespective of treatment, for comparison of primary and secondary counters.

The same prescribed characteristics used for manual enumeration of CN (described above) were incorporated into an algorithm to automate counting using Visiopharm image analysis software. Features of artificial intelligence and machine learning capabilities were used on twice the area that included the same ROIs from which manual counts were assessed to improve accuracy and sensitivity (1.25 x 10^5^ μm^2^/each). For validation and the typical diversity of background staining characteristics, snapshots were randomly chosen to reflect a maximum range of low, median, and high CN counts and variability of backgrounds in immunohistochemically stained cervix sections. These same ROIs were used to train the algorithm with 1 x 10^5^ iterations to achieve a stable flat portion of the asymptotic curve. For initial training, the algorithm ‘learned’ based upon size and intensity to create a smooth perimeter around each methyl green-stained object, which was twice the area used for manual counts to more accurately estimate CN density (as normalized to respective areas). Subsequently, ROIs were batch processed by this automated approach and compared to manual estimates of average CN density (Supplemental Fig. 1). Panel a (left) is one half of an ROI that was counted by the manual approach. Panel b (right) represents the automated counts as determined with the Visiopharm algorithm. Blue dots indicate the vast majority of CN identified by both approaches. Red dots indicate the few discrepant false negative or false positive CN that were misidentified by the Visiopharm algorithm versus manual counts. White dots indicate manual counts of cell nuclei in F4/80-stained macrophages, all of which were included in both CN counting approaches. As described below, the automated CN counts algorithm was used on the same ROI (Supplemental Fig. 1). Comparison of manual versus Visiopharm approaches in this portion of the ROI identified CN (195 and 197, respectively).

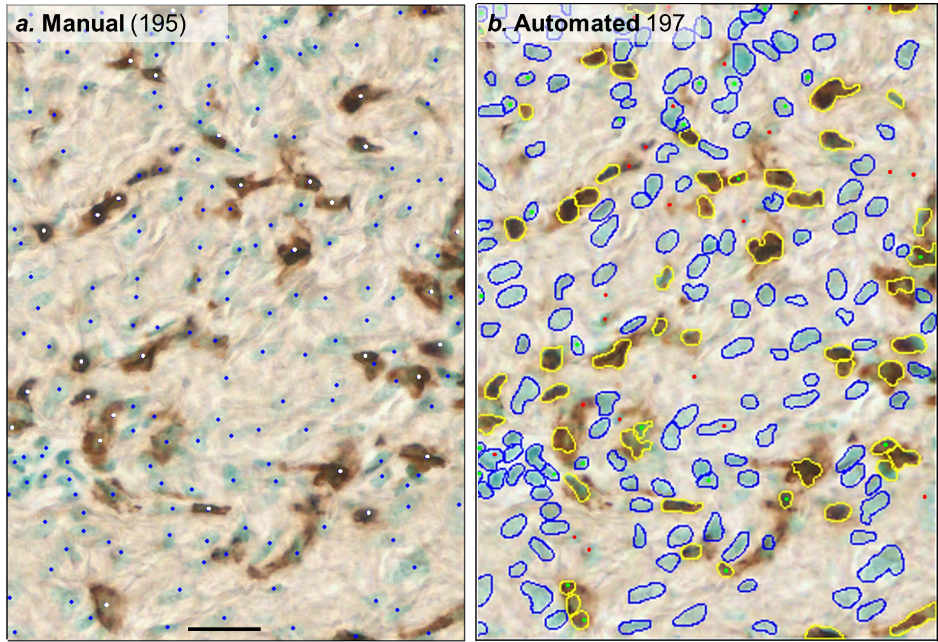
**Supplement Fig. 1** Panel a. Photomicrograph of the ectocervix stroma in an ROI that was used for manual counting of cell nuclei (CN, blue dots) of presumed fibroblasts and CN in DAB-stained macrophages (white dots). Scale bar is 25 μm.

Panel b. The Automated Visiopharm algorithm identified CN (blue perimeter) and macrophages (yellow perimeter).

For validation of the Automated counts approach, the same ROIs used for manual counts were batch processed using the Visiopharm image analysis algorithm. Comparison of automated counts using the Visiopharm and manual approaches produced comparable CN counts (p=0.314, df=42; Supplemental Table 1). Though not significantly different, the small consistently higher counts in automated versus manual approaches were perceived to result from inclusion of CN associated with vascular tissue in the stroma and improved coefficient of variation with automated versus manual approaches, 14% vs 22%. This variability reflects the maximum ‘assay’ range for this unique validation dataset. Furthermore, a sampling of 122 ROIs in the 22 mice in this validation dataset produced an overall discrepancy within an average of ± 9%.

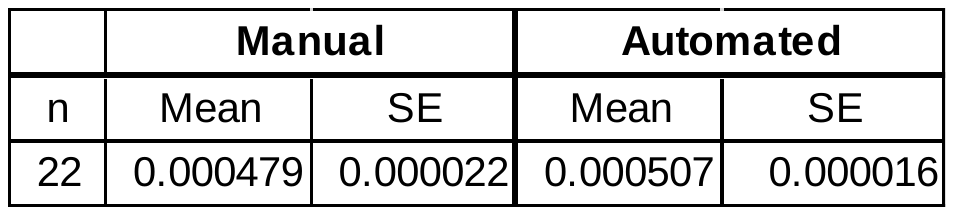
**Supplement Table 1** Mean ± SE of cell nuclei (CN) counts using manual and automated approaches to image analyses of stromal regions in the ectocervix and endocervix from prepartum and postpartum mice (n/group). The protocol for manual counts that is detailed above and summarized in Methods was the same as previously reported.
